# Passage of transmissible cancers in the Tasmanian devil is due to a dominant, shared peptide motif and a limited repertoire of MHC-I allotypes

**DOI:** 10.1101/2020.07.03.184416

**Authors:** A Gastaldello, SH Ramarathinam, A Bailey, R Owen, S Turner, A Kontouli, T Elliott, P Skipp, AW Purcell, HV Siddle

## Abstract

Transmissible cancers are spread via the passage of malignant cells. The survival of the Tasmanian devil, the largest marsupial carnivore, is threatened by two independent transmissible cancers, devil facial tumour (DFT) 1 and devil facial tumour 2 (DFT2). To aid the development of a peptide vaccine and to interrogate how histocompatibility barriers can be overcome, we analysed the peptides bound to Major Histocompatibility Complex class I molecules from the Tasmanian devil and its transmissible tumours. Comparison of the peptidomes from DFT1+IFNγ, DFT2 and host fibroblast cells demonstrates a shared motif, despite differences in MHC-I allotypes between the cell lines. Importantly, DFT1+IFNγ and DFT2 share the presentation of peptides derived from neural proteins, reflecting a common cellular origin that should be exploited for vaccine design. These results suggest that some polymorphisms between tumours and host are ‘hidden’ by a common peptide motif, providing the potential for permissive passage of infectious cells.

## Introduction

Contagious cancers are clonal cell lines that can spread within a population through the passage of live cells between individuals. The emergence in nature of these types of cancers is rare, with only nine described so far, three in mammals and six independent transmissible cancers found in bivalve molluscs (2, 3). The mammalian transmissible cancers include the canine transmissible venereal tumour (CTVT) and two genetically independent transmissible cancers in the Tasmanian devil, devil facial tumour 1 (DFT1) and devil facial tumour 2 (DFT2) (4, 5). Tasmanian devils are the largest surviving carnivorous marsupials found only on the Australian island of Tasmania, but since the emergence of DFT1 (6) their number has declined by 80% (7). The discovery in 2014 of a second independent transmissible cancer, termed DFT2, has the potential to dramatically increase the pressure on population survival (4, 8). Although DFT2 has not been studied extensively, both cancers are thought to spread through biting behaviour, common in devils during feeding and mating, and cause death by starvation and/or organ failure within six to twelve months (5, 9).

DFT1 and DFT2 are effectively allogeneic transplants and should be rejected by the host’s immune system through the engagement of classical major histocompatibility complex class I (MHC-I) molecules on the cell surface of cancer cells with the T cell receptor (TCR) of CD8^+^-T cells. Classical MHC-I molecules are found on all nucleated cells and are responsible for presenting intra-cellular peptides to CD8^+^-T cells. In so doing these molecules define the immunological “self” of an organism. In the presence of viral or cancer derived abnormal proteins the repertoire of peptides presented will be recognised as “non-self” and a cytotoxic response activated by T cells (10). Classical MHC-I molecules are the most polymorphic genes in vertebrates, while non-classical molecules possess more restricted polymorphism and expression as well as having a range of functions (11). Mature cell surface MHC-I molecules are a trimer, composed of a heavy chain, β2 microglobulin (β2m) and an antigenic peptide, usually 8–12 amino acids long, which binds in the Peptide Binding Region (PBR) formed by the α1 and α2 domains of the heavy chain. Peptide binding is an iterative process that occurs in the Endoplasmic Reticulum (ER), where a number of chaperone proteins interact with the MHC-I and β2m to ensure that peptides with high specificity for a given MHC-I molecule are loaded and thus presented to T cells (10).

The amino acids that define whether a peptide will bind to a MHC-I molecule are termed anchor residues, and in most eutherian mammals are found at position two (p2) and the C-terminus (pΩ) of the peptide sequence (12, 13). The binding specificity of an MHC-I allele is determined by polymorphic amino acids that line the PBR and biochemically define a number of pockets that govern interaction with the anchor residues of the peptide ligand. This means that each MHC-I allele has a preferred binding motif that generally describes the salient sequence features of the bound peptide ligands.

In devils, three polymorphic classical MHC-I genes (Saha-UA, -UB and -UC) and three non-classical genes (Saha-UD, -UK and -UM) have been identified, however very little is known about their function or the nature of their ligands (14). Indeed, outside of well characterised species such as human and mouse and those of agricultural importance there has been little analysis of MHC peptide binding in wild species (with a notable exception of the bat MHC (15–17)).

DFT1 cells express low levels of MHC-I, explaining the lack of rejection by CD8^+^-T cells (18). MHC-I down-regulation in these tumours is due to epigenetic mechanisms and can be reversed by treatment with the pro-inflammatory cytokine interferon gamma (IFNγ) (18). However, rare instances of spontaneous DFT1 tumour regression have been described and vaccination and immunotherapy of devils with live MHC-I positive DFT1 cells demonstrate that devils can mount an effective immune response against the disease in some instances (19–21). We have recently shown that in contrast to DFT1, DFT2 cells express MHC-I molecules, but the expressed alleles are similar between hosts and tumours in the samples analysed (1).

In recent years, an increased interest in the discovery of patient-specific tumour neo-antigens has enabled the development of robust experimental work-flows for defining MHC-I peptides in human (22). Here we have adapted and applied these techniques to define the repertoire of peptides (the immunopeptidome) presented by MHC-I molecules in DFT1+IFNγ, DFT2 and their host. Our data suggest that the peptide binding motif of MHC-I molecules on DFT1+IFNγ and DFT2 cells is similar to that of host cells, with a dominant peptide motif characterised by p3 and pΩ anchors. This is the first characterisation of the immunopeptidome of a wild species and its contagious cancers, representing a milestone for the understanding of the immunological mechanisms driving cancer transmission among individuals. In addition, this work will aid the development of prophylactic peptide vaccines to help the conservation of this iconic species.

## Results

### DFT1+IFNγ and DFT2 cells have a restricted repertoire of MHC-I alleles compared to Fibroblasts

Analysis of the MHC-I allotypes on each cell line showed that DFT1+IFNγ and DFT2 cells have a more restricted repertoire compared to fibroblasts, with the majority of allotypes shared between the cell lines (shaded boxes in figure 1a and supplementary table 1).

**Figure 1.**
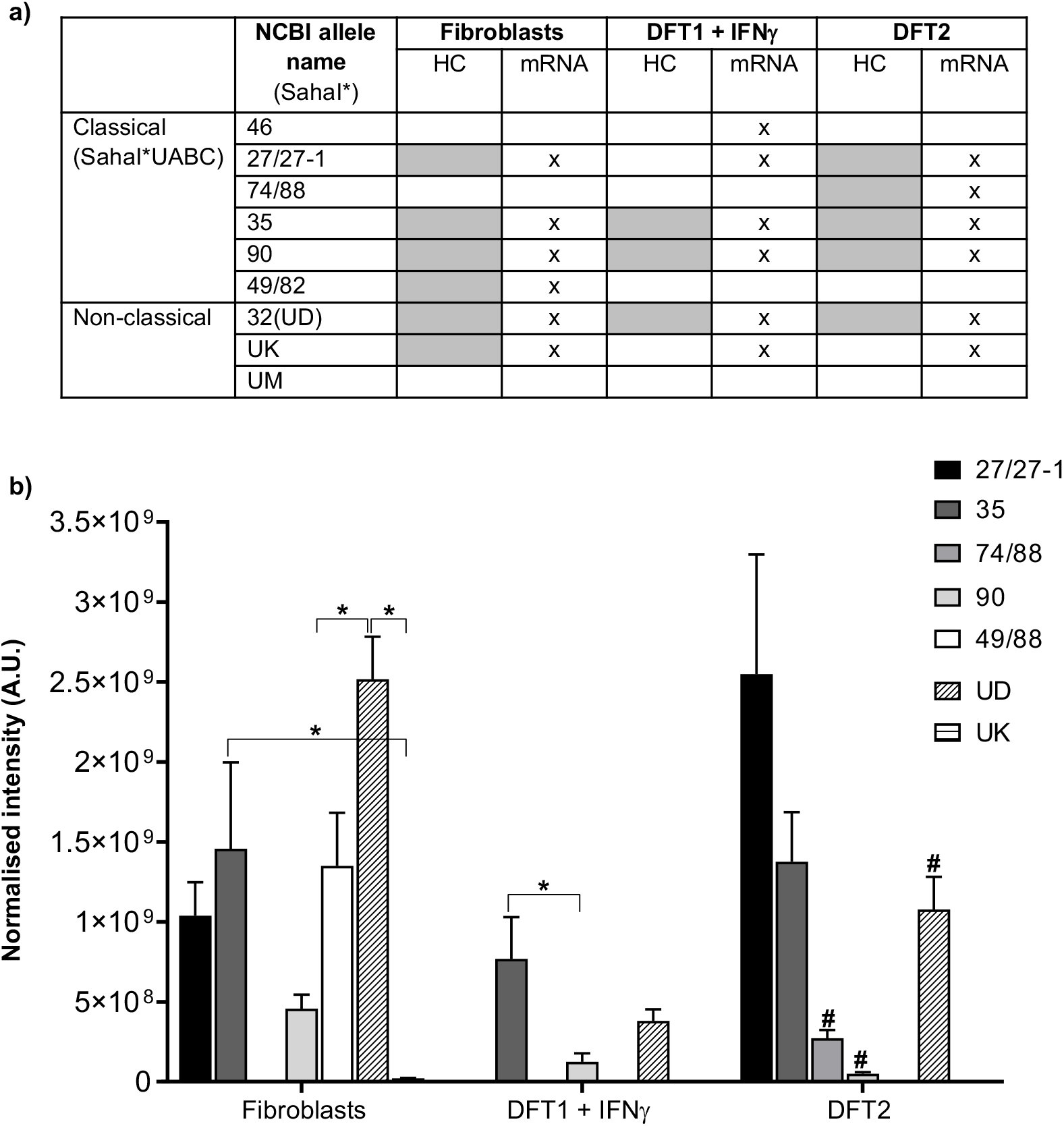
DFT1+IFNγ and DFT2 cell lines have a restricted repertoire of MHC class I (MHC-I) alleles compared to devil fibroblasts. (a) Table of devil MHC-I alleles identified in at least three out of four replicates in devil cell lines by analysis of the heavy chain (HC) fraction separated by HPLC in this paper (shaded boxes) or by mRNA analysis (crossed boxes) in a previous publication (1). A devil fibroblast cell line (referred to as Fibroblasts) represents a host devil. DFT1 is represented by the cell line 4906 treated with interferon γ to induce MHC-I expression (DFT1+IFNγ in the table & text) and DFT2 is represented by the cell line Red Velvet (DFT2 in the table & text). (b) Quantification of HCs of MHC-I alleles in (a) represented as normalised intensity. Statistically significant comparison (p<0.05) are shown with * & #; # = vs SahaI*27/27-1 in DFT2.

These data suggest the Fibroblast cell line has the greatest diversity of cell surface MHC-I molecules with six allotypes in total (SahaI*27/27-1, SahaI*35, SahaI*90, SahaI*49/82, SahaI*32(UD) and SahaI*UK); DFT1+IFNγ cells have the lowest diversity with three (SahaI*35, SahaI*90 and SahaI*32(UD), whilst DFT2 cells have five variants (SahaI*27/27-1, SahaI*74/88, SahaI*35, SahaI*90 and SahaI*32(UD)). Importantly, when DFT1+IFNγ and DFT2 are considered together there is only one unique allotype compared to Fibroblasts, SahaI*74/88 found in DFT2 cells. DFT1+IFNγ cells have no unique allotypes when compared to Fibroblasts. SahaI*27 and SahaI*27-1 were grouped together as the latter differs from SahaI*27 by one non-synonymous substitution in the α2 domain of the heavy chain and the two could not be distinguished based on the peptides analysed by MS. For Fibroblasts, the data mirrored those obtained with mRNA analysis, but in DFT1+IFNγ and DFT2 whilst all the MHC-I variants found at protein level were also found at mRNA level, not all the alleles present in mRNA were consistently identified in the protein pool.

Analysis of the relative abundance of each MHC-I allotype using the intensity of proteotypic peptides found in at least three replicates showed that the allotypes vary in abundance between the cell lines. For SahaI*35, SahaI*49/82, SahaI*32(UD) and SahaI*UK, two unique peptides were quantified that matched each allele, so mean abundance for these variants were calculated (supplementary table 1 for values). The non-classical MHC-I SahaI*32(UD) is the most abundant MHC-I in Fibroblasts lysates (figure 1b), whilst SahaI*27/27-1 was the most abundant in DFT2 and SahaI*35 and SahaI*32(UD) were the most abundant variants in DFT2. The only allotype found in DFT2 and not in Fibroblasts, SahaI*74/88, had a relatively low abundance. Saha*UK was only present in Fibroblasts and was the least abundant MHC-I allotype. Thus, SahaI*27/27-1, SahaI*35 and the non-classical SahaI*32(UD) are the most consistently expressed across the cell lines, but there are differences in allotyopes and their level of expression.

### Tasmanian devil MHC-I molecules preferentially bind peptides of 8 and 9 amino acids in length

We used immunoaffinity purification with a devil specific anti-β2m antibody to isolate the peptides associated with the MHC-I heavy chains identified in DFT1+IFNγ, DFT2 and fibroblast cell lines. In total, from all three cell lines (n=4), we isolated 25,554 unique peptides that were 7-15 amino acids, the expected length preference for MHC-I molecules (7-15mers, table 1).

**Table 1.**
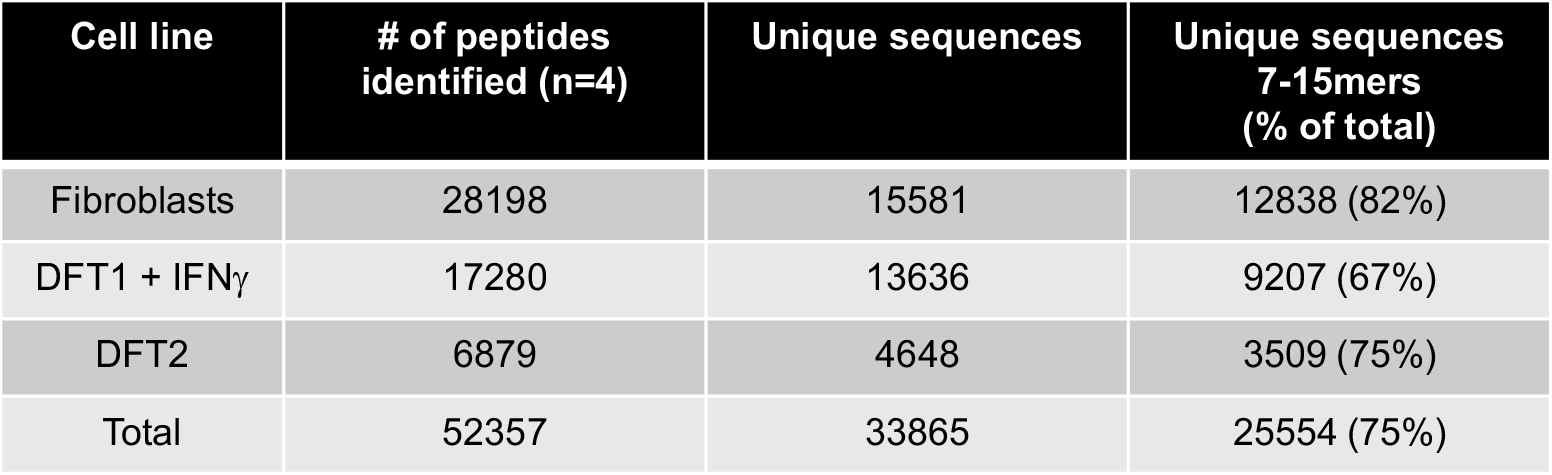
Number of total and unique peptide sequences in a devil fibroblast cell line and in devil facial tumour 1 and 2 (DFT1+IFNγ & DFT2) cell lines. Peptides were isolated by immunoaffinity purification with a pan-MHC-I antibody against devil β2m followed by separation with HPLC and analysis by mass spectrometry (n=4/cell line). The fibroblast cell line (Fibroblasts) yielded the highest number of unique 7-15mer peptide sequences (12,838) across the 4 replicates, with lower numbers isolated from DFT1+IFNγ (9,207) and DFT2 (3,509). The number of peptides identified correlates with cell surface β2m expression on each cell line, with Fibroblasts showing the highest expression, DFT2 the lowest and DFT1+IFNγ intermediate (supplementary figure 1).

Analysis of the length distribution of unique peptides (figure 2a) revealed that 8mers were more abundant than 9mers in Fibroblasts and DFT2 cells but not in DFT1+IFNγ cells where 9mers were most abundant. Shorter and longer peptides were identified with lower frequency, and while similar proportions of 10mers were found in all cell lines (figure 2a, % unique sequences = Fibroblasts, 8.4%; DFT1+IFNγ, 8.1%; DFT2, 8.4%), DFT1+IFNγ and DFT2 cell lines displayed higher proportions of longer peptides in comparison with Fibroblasts (supplementary table 2).

**Figure 2.**
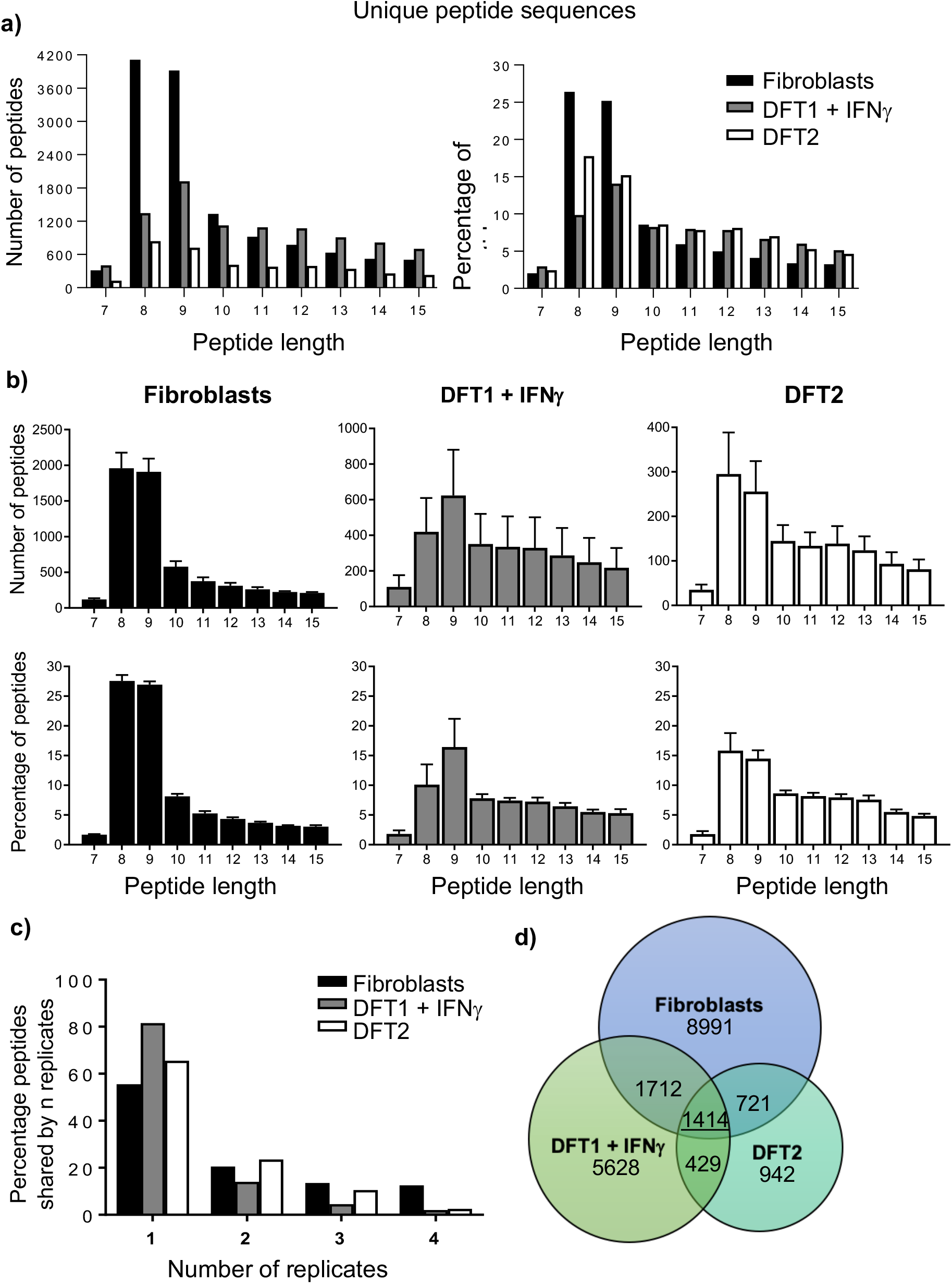
Features of peptides from devil cell lines. (a) Length distribution of unique peptide sequences across all replicates isolated from Fibroblasts, DFT1+IFNγ and DFT2 cells. (b) Total number and percentage of peptides of different lengths. Data are mean ± SEM, n=4/cell line. (c) Graph of the percentage of peptides found in “n” replicates for each cell line as a measurement of experimental reproducibility. (d) Venn diagram showing unique 7-15mer peptides shared among Fibroblasts, DFT1+IFNγ and DFT2.

We calculated the percentage of 7-15mers found in either one, two, three or four replicates across all three cell lines. This analysis showed that the greatest proportion of peptides was found in only one replicate for all three cell lines, with DFT1+IFNγ displaying the highest single replicate bias (figure 2c). The three cell lines shared 1414 unique 7-15mers (figure 2d & supplementary table 4), whilst DFT1+IFNγ /Fibroblasts, DFT2/Fibroblasts and DFT1+IFNγ/DFT2 shared respectively 1712, 721 and 429 unique sequences 7-15 amino acids in length.

### Peptides from Fibroblasts, DFT1+IFNγ and DFT2 possess similar anchor residues found at pΩ and p3 sites

The consensus-binding motifs of peptides isolated from Fibroblasts, DFT1+IFNγ and DFT2 show striking similarities, despite the differences in MHC-I allotypes between cell lines. The amino acid frequencies were calculated for each amino acid at each position in 8mers and 9mers (supplementary table 4) and classified as dominant if present with a frequency >30%, strong if found with a frequency 20-30%, and moderate if observed at 10-20% (figure 3).

**Figure 3.**
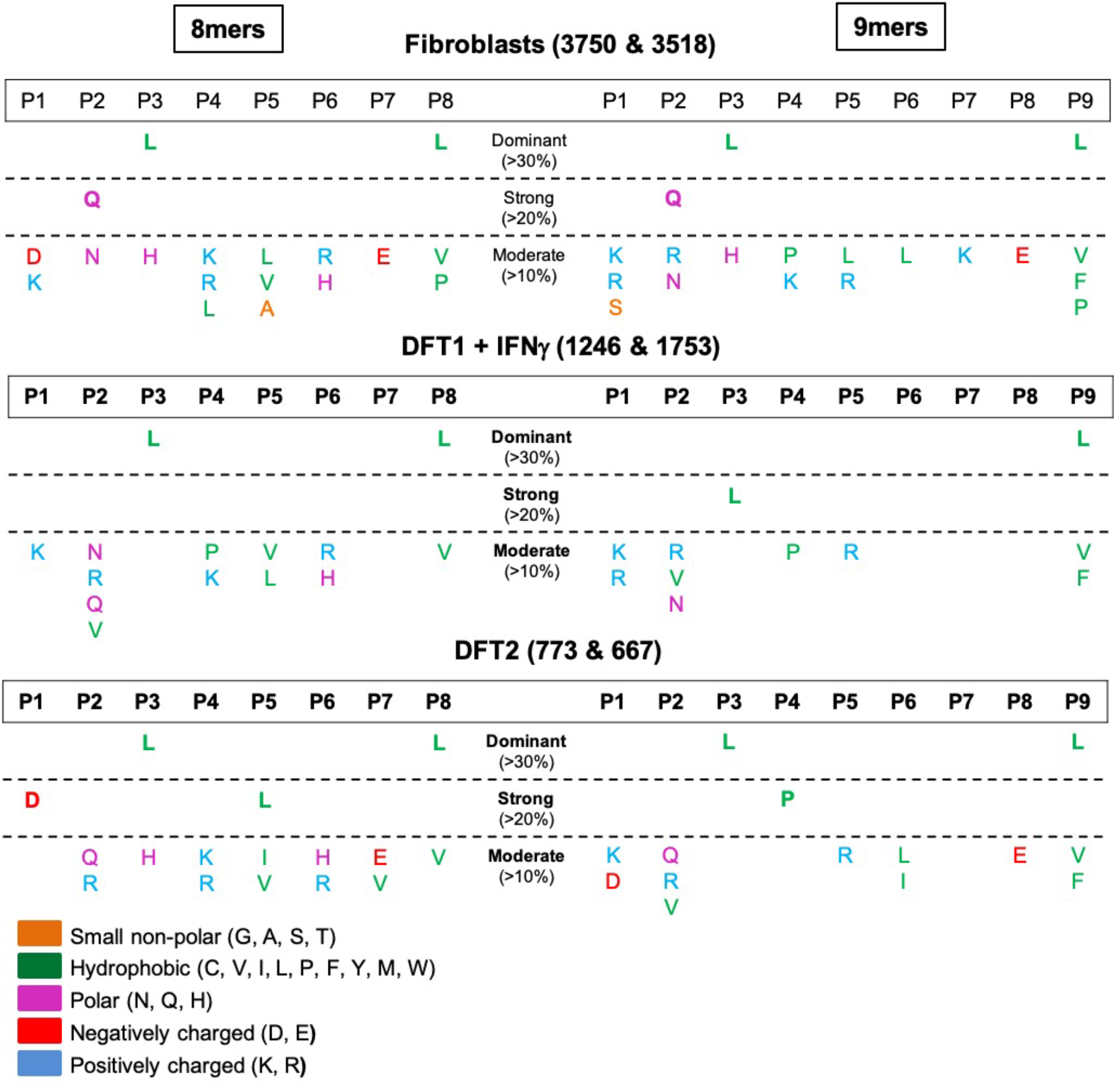
DFT1+IFNγ, DFT2 and Fibroblasts share a preference for leucine anchor residues at p3 and pΩ. Tables of consensus binding motifs of 8mer and 9mer peptides in devil cell lines. Motifs were derived by calculating the frequency of each single amino acid at each position. Numbers after cell line names refer to the number of unique 8mers & 9mers analysed across replicates (n=4). P1-P9 = positions in the peptide sequence. Amino acids are represented by their single letter abbreviations, and are coloured according to their biophysical properties based on the LESK scheme.

The 8mer and 9mer peptides had a preference for the hydrophobic amino acids leucine (L), valine (V), phenylalanine (F) or proline (P) at the C-terminus (position Ω, pΩ) in all three cell lines (figure 3), a characteristic common to many eutherian MHC-I peptides. All three cell lines also have a preference for L at p3 in both 8mers and 9mers, with this being dominant in Fibroblasts and DFT2 and strong in DFT1+IFNγ. Alongside this dominant motif there are also features specific to each cell line. Glutamine (Q) was strong at p2 in 8mers and 9mers from Fibroblasts, while aspartic acid (D) was strong at p1 and L at p5 for DFT2 8mers and, P was strong at p4 in 9mers from DFT2.

To further validate the consensus motif, additional immunopeptidome experiments were performed on a smaller number of DFT2 cells (1×10^8^/replicate). 4450 peptides were isolated across three replicates, and the distribution of both the number and percentage of total 7mers to 15mers (supplementary figure 2a and supplementary table 5) were similar to that of peptides from previous experiments using 1×10^9^ cells/replicate of DFT2 cells (figure 2a). Consensus binding motifs (supplementary figure 2b) supported the finding of a dominant motif with hydrophobic anchors at pΩ (L or F) and p3 (L) (figure 3) and potential additional anchors at p1 and p5 in 8mers, and at p4 in 9mers.

To complement these analyses, we used the GibbsCluster algorithm (23) to analyse the immunopeptidome dataset and cluster the peptides into similarity groups based on peptide sequence features (supplementary figure 3a-b). This analysis returned the highest probability (represented by the “Kullbach Leibler Distance (KLD) for the peptides deriving from a single group (cluster) with a motif of L at p3 and L at pΩ for all three cell lines. This was regardless of whether the analysis was directed by the number of MHC-I allotypes found in each cell line. However, it is difficult to distinguish a single group from the likelihood of peptides deriving from up to 6 different groups, indicating that peptide groups with a less dominant motif might also have been present in the data set. This is consistent with our finding that the peptides derived from between 3 and 6 MHC-I alleles present on the cell lines and that the alleles vary in abundance.

### Devil MHC class I F-pocket, but not B-pocket, shows homology with MHC class I molecules from eutherian mammals

In the absence of structural data for devil MHC-I molecules, homology modelling and intraspecies pairwise sequence comparisons reveals differences in the classical binding pockets of the devil MHC-I that are consistent with the peptide motifs observed above. Ten devil MHC-I alleles were modelled against the recently described bat (*Pteropus Alecto*) Ptal-N*01:01 structure (15) to predict both the B-pocket (accommodating p2) and F-pocket (accommodating pΩ) conformations. The bat Ptal-N*01:01 allele was used for the analysis as it shares a 3aa insertion in the α1 domain with devil MHC-I when compared to other eutherian MHC-I alleles and therefore could provide a more matched homology model. Conformation-weighted pairwise comparisons were made against 16,748 MHC-I sequences from seven species (predominantly human). Only two devil MHC-I alleles showed a high degree of conservation in their sequence across the predicted B pocket: SahaI*27 and SahaI*74/88 (figure 4a and figure 4c). Both alleles were identical to HLA-B*08:04 at the residues making up the B-pocket (figure 4c, red shading) resulting in the same predictions. Across the F-pocket (figure 4a, righthand side), seven devil MHC-I alleles showed high conservation in their sequence, four of which were identical predictions due to identical F-pocket sequences (figure 4c, blue shading). Only SahaI*90 and SahaI*UK alleles did not have a predicted F pocket. A summary of binding pockets and corresponding epitope predictions is given in figure 4b. Predicted anchor residues at the B-pockets of SahaI*27 and SahaI*74/88, but an absence of homology at the B pocket for the remaining alleles, is consistent with the experiementally derived consensus binding motifs that predict only a secondary anchor at p1 or p2. Further, the high similarity of the F pocket supports the presence of a hydrophobic pΩ anchor. Interestingly, while the F pocket of SahaI*32(UD) supports the presence of a hydrophobic anchor at pΩ this was not be predicted for SahaI*UK suggesting this non-classical molecule might have not contributed to the immunopeptidome. Indeed, whilst SahaI*32(UD) was found to be present at high levels in the heavy chain fraction analysis, SahaI*UK was present at very low level only in Fibroblasts (figure 1).

**Figure 4.**
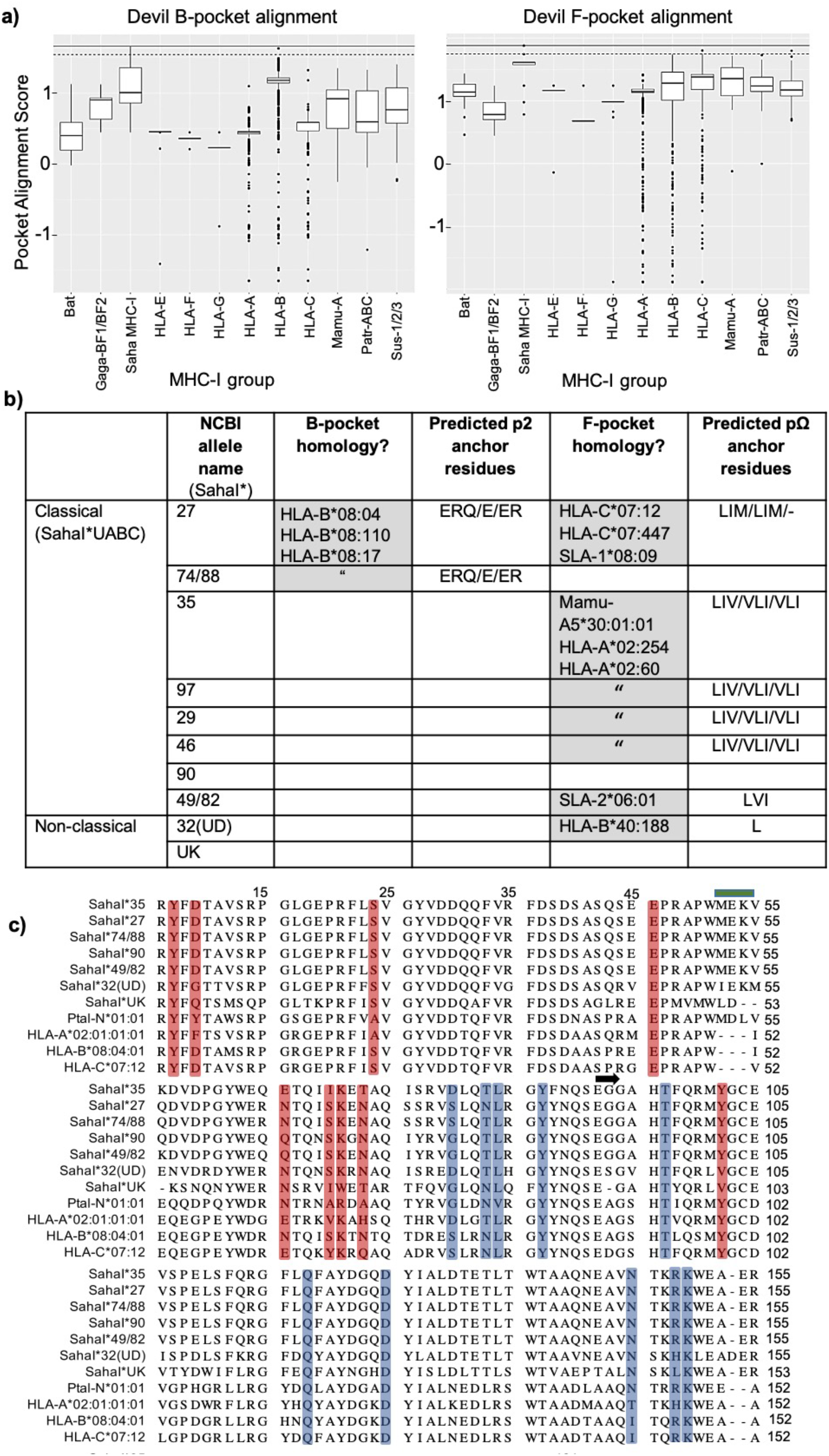
The F pocket of devil MHC-I alleles is conserved with MHC-I from eutherian mammals. (a) Binding pocket homology analysis of devil MHC-I allele SahaI*27 against 16,748 MHC-I sequences across 12 different MHC-I supertypes from different species: Gaga = *Gallus gallus*, Mamu = *Macaca mulatta*, Patr = *Pan troglodytes*, Sus = *Sus scrofa*. Upper black line shows perfect alignment score, datapoints above the tapered cut-off line indicate sequence homology and were used to predict pocket motifs. (b) Table showing where homology was identified for B-pocket and/or F-pocket (grey boxes) for each devil MHC-I allele; if predictions were possible, the respective motifs of the top 3 matches within that pocket have been included (p2/b-pocket, pΩ/F-pocket). The alleles with the highest homology to the devil are reported inside each grey box. (c) Multiple sequence alignment of α1-α2 peptide binding groove region of devil MHC-I alleles with selected human HLA alleles and the Ptal-N*01:01 bat MHC-I allele. HLA-B*08 and HLA-C*07 were included as they had the highest alignment scores to SahaI*27 with the B and F pocket respectively. B-Pocket residues are highlighted in red and F-Pocket residues are highlighted in blue. The blue bar denotes the 3aa insertion common to Ptal and devil MHC-I sequences. A black arrow marks the beginning of the α2 domain.

### Dominance of leucine (L) at p3 of 8mers and 9mers is a unique characteristic of devil peptides

The amino acid leucine is prevalent at pΩ in many human and murine MHC-I alleles, while dominance of L at p3 is less common. We compared the frequency of L at p3 in devil peptides to a range of well characterised MHC-I allotypes found in human and mouse (supplementary table 6). As seen in figure 5a, leucine was much more frequent at p3 in 8mers and 9mers isolated from devil cell lines than among peptides recognised by a variety of human and murine alleles. The alleles with the highest frequency of L at p3 among 9mers were the murine H-2 Db and H-2 Kd, although they did not reach frequencies found in devil. The percentages of small and bulky amino acids present in devil MHC-I bound peptides were also calculated (figure 5b). As reported previously for other species (17, 24–26), the use of small amino acids increased whereas the frequency of bulky amino acids decreased with increasing peptide length in devil (supplementary table 6).

**Figure 5.**
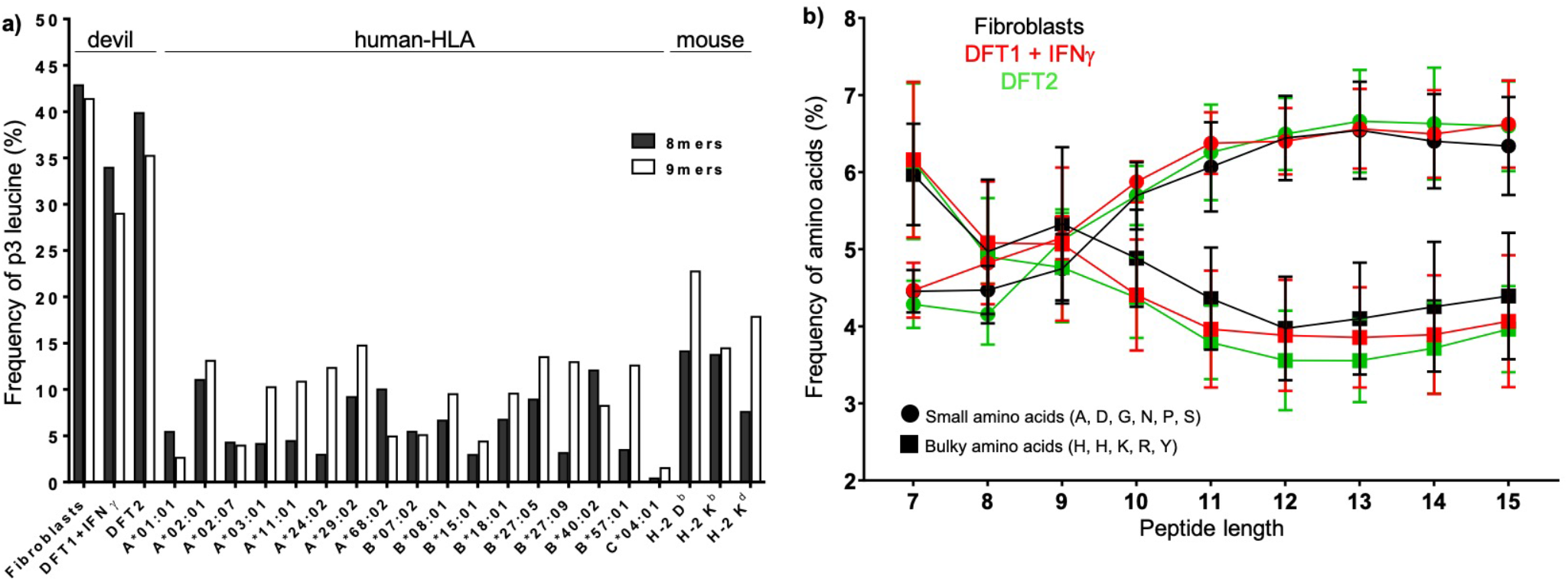
The strong preference for the amino acid leucine (L) at p3 in 8mers and 9mers is a unique characteristic of devil cell lines, whilst small versus bulky amino acid usage is similar to that of other species. (a) Observed frequency (%) of L at p3 in 8mer and 9mer peptides from devil cell lines (n=4) compared with those calculated for selected mammalian alleles found in human and mouse. (b) Frequency of small and bulky amino acid usage was determined for each peptide length in the range 7-15 amino acids. Alanine (A), aspartate (D), glycine (G), asparagine (N), proline (P) and serine (S) constituted small amino acids while phenylalanine (F), histidine (H), lysine (K), arginine (R) and tyrosine (Y) were considered bulky amino acids. Data are mean ± SEM.

### Peptides derived from neural specific source proteins are presented on MHC-I molecules in DFT1+IFNγ and DFT2

Annotations for source proteins of peptides in Fibroblasts, DFT1+IFNγ & DFT2 were retrieved and compared with those from three human MHC-I alleles, two mouse alleles and one bat allele (figure 6 & supplementary table 7 for full results). The greatest percentage of peptides were derived from either nuclear or cytoplasmatic proteins (figure 6a) and involved in regulation and cellular processes (figure 6b), for all analysed cells/alleles. GO slim name (supplementary figure 4a-b & supplementary table 7 for full results) analysis confirmed that the majority of peptides had originated from cytoplasmatic and nuclear proteins involved in biosynthetic process, cellular nitrogen compound metabolic processes, anatomical structure development and transport.

**Figure 6.**
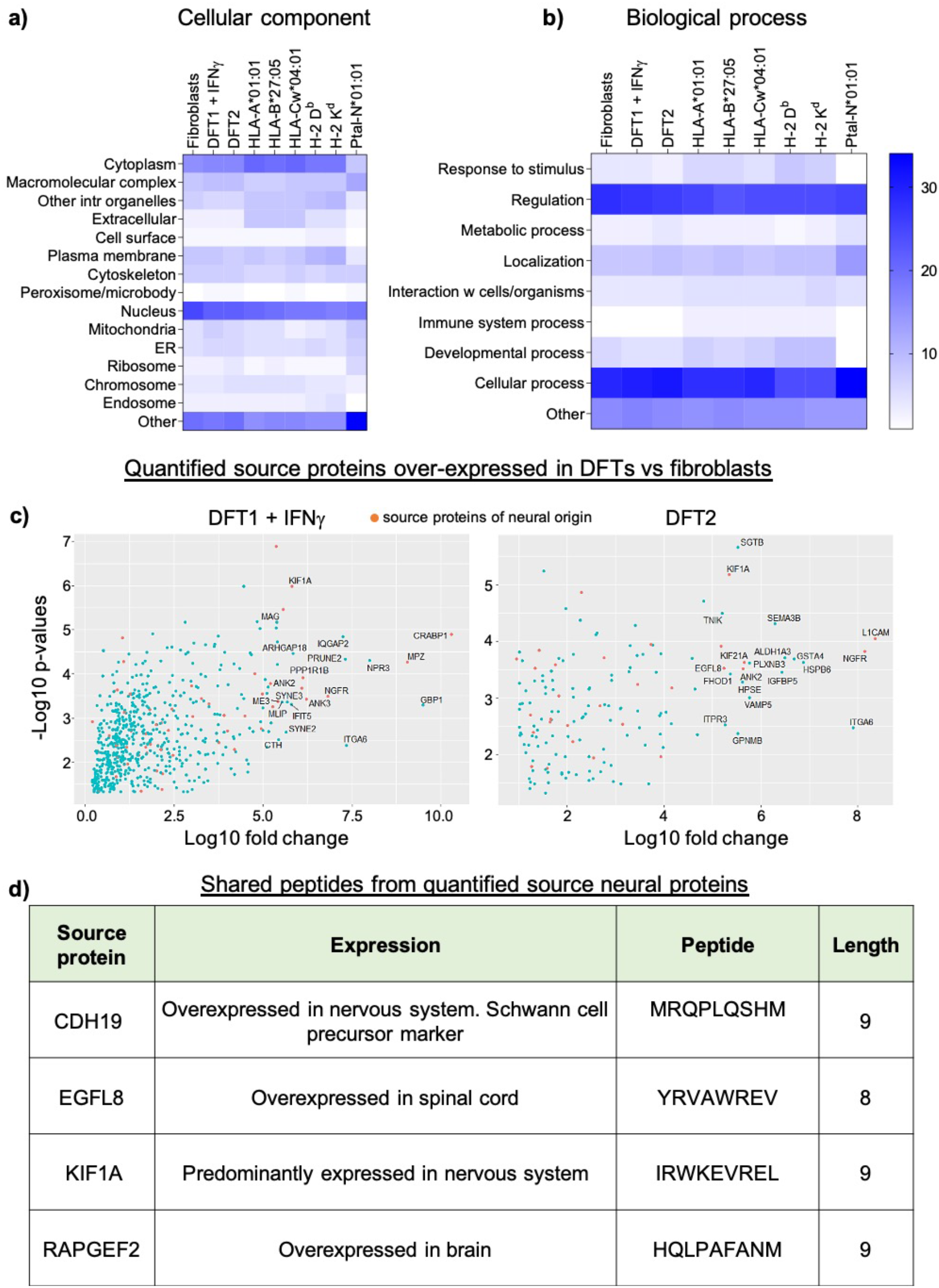
Neuronal proteins are a source of shared peptides in DFT1+IFNγ and DFT2 cell lines. Heat maps representing the percentage of source proteins from different cellular components (a) or biological processes (b) for peptides isolated from Fibroblasts, DFT1+IFNγ and DFT2 compared with three MHC class I alleles found in human (HLA-A*01:01, HLA-B*27:05, HLA-Cw*04:01), two found in mice (H-2 D^b^, H-2 K^d^) and one found in Pteropus alecto (the Australian black flying fox, a bat species (Ptal-N*01:01)). Percentages are represented on a sliding scale of the colour blue, were higher percentages corresponds to higher intensity of blue. (c) Volcano plots representing quantified proteins, which are a source of peptides and over-expressed in DFT1+IFNγ (left) and DFT2 (right) compared to host fibroblasts. Proteins whose expression is restricted to or more abundant in the nervous system (neural proteins) are coloured in orange; proteins with the highest expression are labelled. (d) Table listing the neural source proteins of 8mer and 9mer peptides found in both DFT1+IFNγ and DFT2 cell lines but not in Fibroblasts. Expression pattern and peptide sequences are also given.

As DFT1+IFNγ and DFT2 are are predicted to have the same cellular origin, a Schwann cell (27–29), we identififed the peptides that derive from source proteins with a neural function that are overexpressed in DFT1+IFNγ and/or DFT2 compared to Fibroblasts. Quantitative proteomic analysis of devil Fibroblasts, DFT1+IFNγ and DFT2 (datasets in supplementary table 8) was used to identify proteins normally specific to, or overexpressed in the nervous system (referred to hereon as neural proteins) that were over-expressed in DFT cell lines relative to Fibroblasts and a source of ligands for the devil MHC-I allotypes (figure 6c and supplementary table 9). In DFT1+IFNγ, 54 peptides (8mers and 9mers) were derived from 45 neural proteins which were not a source of peptides in Fibroblasts. In DFT2, 15 peptides (8mers and 9mers) were identified that derived from 17 neural source proteins and did not appear in the Fibroblast immunopeptidome (supplementary table 9). Four of these source proteins were shared between DFT1+IFNγ and DFT2 and generated 1 shared 8mer (source protein: EGFL8) and 3 shared 9mers (source proteins: CDH19, KIF1A, RAPGEF2) (figure 6d). Among the shared proteins we identified the Schwann cell precursor marker CDH19, and among the twenty highest expressed neural source proteins found in DFT1+IFNγ but not in DFT2, we identified the Schwann cell markers Myelin-Associated Glycoprotein (MAG) and Myelin Protein Zero (MPZ). Among those unique to DFT2 we found the early glial markers Nerve Growth Factor Receptor (NGFR).

## Discussion

This is the first study to identify endogenous peptides from the MHC-I molecules of a marsupial species and our analysis joins only a limited number of studies on MHC-I derived peptides from wild species (14, 16). Here, we analysed the repertoire of peptides bound by MHC-I molecules (the immunopeptidome) of Tasmanian devil fibroblasts and the transmissible cancer cell lines, DFT1+IFNγ and DFT2, to define the antigenic landscape of the tumour cells and to provide an evidence based approach for vaccine design. We show that DFT1+IFNγ and DFT2 cells have a more limited repertoire of MHC-I molecules compared to host cells. Comparing the peptides isolated from the tumour and host cells lines reveals a similar, dominant motif with a preference for Leucine (L) at p3 and pΩ. The shared motif between the cell lines can not be fully explained by shared MHC-I allotypes, hence different devil MHC-I allotypes may share a preferred peptide motif. The restricted MHC-I repertoire and dominant peptide motif reduces the antigenic variation on DFT1+IFNγ and DFT2 compared to host, potentially allowing the tumour cells to mimic the host.

The DFT1+IFNγ, DFT2 and fibroblast cell lines share a preference for L at p3 and L or another hydrophobic residue at pΩ. The presence of an anchor residue at pΩ, and a preference at this position for hydrophobic amino acids, has also been observed in many mammalian and nonmammalian MHC-I alleles (12, 30, 31). Less common among other species is the presence of an anchor residue at p3 (with a weaker anchor at p2). Notable exceptions are HLA-A*01:01, with an acidic E/D at p3 (25) and HLA-C*04:01, with a preference for D at p3 and a secondary anchor at p2 (12, 32). Our analysis of >16,000 MHC-I alleles and homology modelling of the devil MHC-I allele SahaI*27 against the recently described bat MHC-I structure (Ptal-N*01-01) (15), shows a high level of conservation within the F pocket that is absent in the B pocket and likely explains the different binding properties of the devil MHC-I molecules. Interestingly, analysis of all MHC-I alleles expressed by the tumour and host cell lines predicts a B-pocket preference for only SahaI*27 and SahaI*74/88 (which differ by only one amino acid across the α1 and α2 domains), but a F-pocket prediction for a hydrophobic residue is largely consistent across the alleles. It is possible that certain devil MHC-I molecules bind peptides through the B-pocket and utilise a p2 anchor or a p1 anchor. Given our analysis used a pan specific anti-β2m antibody we have likely captured multiple peptide motifs, some of which are partially ‘hidden’ in our dataset due to differences in expression level among MHC-I allotypes. Using the data generated here it should be possible to determine the exact binding properties of particular MHC-I allotypes using structural analyses.

The use of the anti-β2m antibody for immunoaffinity purification of MHC-I molecules presents a problem for assigning specific peptide motifs to individual devil MHC-I allotypes. A Gibbs cluster analysis predicts that the peptides from each cell line form a single cluster (predicting a L at p3 and L at pΩ). However, this prediction is not statistically strong and based on our analysis of the heavy chain fractions isolated from the purification experiments we conclude that the peptide pool was derived from multiple alleles. At a minimum, we predict that fibroblasts have six allotypes at the cell surface and DFT1+IFNγ and DFT2 have 3 and 5 functional allotypes respectively. These alleles correlate well with our previous definition of the MHC-I transcripts expressed by DFT1+IFNγ and DFT2 and fibroblasts (1). However, there are a number of exceptions. First, we found no evidence of SahaI*27/27-1 or SahaI*46 in the heavy chain fraction from DFT1+IFNγ cells, despite the presence of mRNAs for these alleles in DFT1+IFNγ. Second, we find only trace amounts of the non-classical MHC-I, SahaI*UK, in all three cell lines, which is surprising given we have previously found high expression of this gene in the tumour cell lines. This could be because SahaI*UK does not bind peptides as is the case for other non-classical MHC-I molecules (33, 34). Certainly modeling of the devil MHC-I alleles to define the pockets of the PBR shows that the F pocket could not be defined for SahaI*UK, perhaps indicating this allotype has a non-canonical pΩ anchor.

Despite the presence of multiple MHC-I allotypes in the heavy chain fractions we find that the cell lines have a dominant motif (L at p3 and L at pΩ) and evidence of a secondary motif, perhaps derived from less well expressed alleles. Further investigation is needed to determine whether this phenomenon is due to devil MHC-I molecules binding similar peptides or to the expression of the same dominant variant in all cells. However, DFT1+IFNγ and DFT2 cells have a more restricted repertoire of MHC-I alleles compared to fibroblasts and all DFT1+IFNγ and DFT2 MHC-I alleles are found on the fibroblast cells, with the exception of SahaI*74/88. If this is a general feature of DFTs when compared to host devils across Tasmania this will give the tumours an advantage in allograft transmission. Within the transplantation literature there is a growing appreciation of the importance of MHC class I and class II supertypes. These supertypes mean that ‘permissive’ HLA mismatches between donor and host result in graft acceptance due to similarities in the peptide motifs presented and molecular mimicry in the interaction of donor peptide/MHC and host TCR (35, 36). It is possible that several closely related devil MHC-I alleles are the same ‘supertype’ and have highly similar peptide binding properties. Further analysis of a large subset of host animals is needed to test this hypothesis. Mammalian MHC-I molecules generally bind to peptides which are 8 to 10 amino acids in length, although several examples of different length distribution preferences have been described (37, 38). Typically, 9mers are the most abundant peptides for the majority of HLA allotypes (13). We found instead that MHC-I molecules on DFT cells and fibroblast cells bind high numbers of both 8mers and 9mers, with DFT2 and fibroblasts having a preference for 8mers. The preference for 8mers may reflect specific binding properties of the devil MHC-I molecules and the preference for 9mers in DFT1+IFNγ cells may be due to stimulation of these cells with IFNγ, which is known to change the repertoire of peptides presented when the immunoproteosome is triggered (26, 39, 40).

DFT2 is a more recently emerged tumour compared to DFT1, and in contrast to DFT1, this tumour does express MHC-I molecules. Although DFT2 is still only found in a limited area, its prevalence is increasing (8). A unique opportunity exists to limit the spread of this ‘new’ tumour, before it causes the level of infection of DFT1. MHC^+^-DFT1 cells have become the focus of intense research to determine if they are allogenic to host devils and have potential as a whole cell vaccine against DFT1 (19, 21). Preliminary results have shown that MHC^+^-DFT1 cells can stimulate a protective immune response in host devils but the mechanisms are not understood and the results are variable among host devils. A successful peptide vaccine relies on the identification of tumour associated or tumour specific antigens on the surface of tumour cells. We have identified four unique 8/9mer peptide sequences deriving from four source proteins which are present on the surface of both DFT1+IFNγ and DFT2, but not fibroblast cells. These peptides derived from source proteins which are specific to or overexpressed in cells of the nervous system (including Schwann cell development), which supports previous studies proposing a common cellular origin for the tumours (29). Further analysis is needed to characterise the peptide repertoires of more tissues in the Tasmanian devil and to identify mutated peptide antigens, but the identification of unique peptides in DFT1+IFNγ and DFT2 from proteins with differential tissue expression indicates that tumour associated or tumour specific antigens may exist in both DFT1 and DFT2.

Here we have demonstrated that there is a restricted MHC-I allele repertoire on transmissible cancer cells combined with a single dominant peptide binding motif similar to that found in host cells. This data may explain why DFT cells are not being rejected by the devil immune system, despite expression of MHC-I molecules. This is of particular relevance for DFT2 compared to DFT1, as DFT1 is MHC-I negative unless stimulated with IFNγ. Indeed loss of MHC-I molecules by DFT1 suggests that there is a limit to the ability of these tumours to transmit while still expressing MHC molecules. Thus, while the ability of DFT1 and DFT2 to mimic host cells without sharing all MHC-I alleles may explain how multiple transmissible cancers have emerged in the Tasmanian devil, more research is needed to establish the limits of cancer transmission across histocompatibility barriers.

## Material and Methods

### Cell culture and IFNγ treatment

Three Tasmanian devil cell lines were analysed, the devil fibroblast cell line ‘Salem’ (refered to as Fibroblasts in text), the DFT1-derived cell line 4906 and the DFT2-derived cell line RV (red velvet, refered to as DFT2), and are described in (1, 4, 41). To stimulate expression of MHC-I molecules in DFT1, cells were cultured (17 h) in medium containing recombinant devil IFNγ according to previously described protocols (18), and are refered to as DFT1+IFNγ in the text. To purify devil MHC-I molecules by immunoaffinity purification, devil cells were expanded (1×10^9^ cells/replicate, n=4) using Corning^®^ HYPER*Flask*^®^ M cell culture vessels (Sigma-Aldrich), harvested and washed in 1X-PBS before being snap frozen in liquid nitrogen and stored at −80°C. Media for cell growth and IFNγ treatment was batch prepared for consistency. Beta-2-microglobulin (β2m) was used to assess total surface MHC-I expression as previously described (1).

### Isolation of MHC-I bound-peptides by immunoaffinity purification

Peptides bound to MHC-I molecules were isolated from cell lysates by immunoaffinity purification of devil β2m protein, purified by reversed phase-high performance liquid chromatography (RP-HPLC) and identified by liquid chromatography-mass spectrometry/mass spectrometry (LC-MS/MS) as described in (22). Four replicates of DFT1+IFNγ, DFT2 and Fibroblasts cells were used containing 10 pooled fractions per replicate. An experiment on DFT2 cells only (n=3) with lower number of cells (1×10^8^) was used to validate the data.

Anti-devil β2m antibody was purified by fast protein liquid chromatography (FPLC) from supernatant of hybridoma cells (identifier: 13-34-38). All chemicals were from Sigma-Aldrich unless otherwise stated; all wash buffers and solutions were made using MS grade H_2_O and are detailed in (22).

<u>Generation of cell lysate</u>: frozen cell pellets (1×10^9^ cells/replicate, n=4) were ground using a cryogenic mill (Mixer Mill MM 400 (Retsch), 30 Hertz, 1 min x 2) followed by incubation (1 h, 4 °C) with constant rotation in lysis buffer. Lysates were centrifuged (2000*xg*, 10 mins, 4 °C) to remove nuclei, and supernatant was clarified by ultracentrifugation (100000*xg*, 45 mins, 4 °C).

<u>Preparation of immunoaffinity columns</u>: 1.5 mL of protein A agarose resin (Repligen) was washed (PBS, 10 c.v.) before incubation with anti-devil β2m antibody (10 mg in PBS) under constant rotation (4 °C, 1h). The antibody bound-resin was washed with borate buffer, followed by freshly prepared triethanolamine to ensure there were no residual amines interfering with the cross-linking reaction. The antibody was cross-linked by incubating (1 h at RT) with Dimethyl pimelimidate (40 mM in 0.2 M triethanolamine, pH 8.2), and the reaction was terminated by adding ice-cold Tris. Unbound antibody was removed by washing with citrate buffer and the column washed with PBS until pH of flow through was > 7.

<u>Immunoaffinity purification of devil MHC-I molecules</u>: cross-linked columns were transferred to 4 °C and cell lysates were passed through the resin twice. Flow throughs were collected and stored at −80°C for subsequent proteomics analysis. The columns were washed at room temperature. MHC-I molecules were eluted with 10% acetic acid.

<u>Separation of MHC-I eluate by RP-HPLC</u>: eluted peptides were separated from MHC-I heavy chains, β2m and other contaminants using a C18 reversed-phase HPLC column (4.6 mm internal diameter x 10 cm, Chromolith Speed Rod, Merck) running on a mobile phase buffer of buffer A (0.1% trifluoroacetic acid (TFA)) and buffer B (80% acetonitrile/0.1% TFA) on an AKTAmicro^™^ HPLC system (GE Healthcare). The separation was performed with a rapid gradient of buffer A to B (2 to 40% B for 4 mins, 40 to 45% for 4 mins and a rapid 2 min increase to 100% B) and fractions (500 μL) collected on LoBind Eppendorf tubes. Fractions were vacuum dried (40°C), re-suspended in 0.1% fresh formic acid (15 μL), sonicated (10 mins) and stored at −80°C.

<u>LC-MS/MS analysis of peptides</u>: thawed samples were sonicated before addition of indexed retention time (iRT) peptide standards (1 pmole) and centrifugation (16060*xg*, 10 mins) to remove any particulates. Peptides were trapped on a 2 cm Nanoviper PepMap100 trap column at a flow rate of 15 μL/min using a RSLC nano-HPLC. The trap column was then switched inline to an analytical PepMap100 C18 nanocolumn (75 μm x 50 cm, 3 μm 100 Å pore size) at a flow rate of 300 nL/min using an initial gradient of 2.5% to 7.5% buffer B (0.1% formic acid 80% ACN) in buffer A (0.1% formic acid in water) over 1 min followed with a linear gradient from 7.5% to 32.5% buffer B for 58 min followed by a linear increase to 40% buffer B over 5 min and an additional increase up to 99% buffer B over 5 min. Separated peptides were analysed using a Q-Exactive Plus mass spectrometer (ThermoFisher Scientific, Bremen, Germany). Survey full scan MS spectra (m/z 375–1800) were acquired in the Orbitrap with 70,000 resolution (m/z 200) after the accumulation of ions to a 5 × 10^5^ target value with a maximum injection time of 120 ms. The 12 most intense multiply charged ions (z ≥ 2) were sequentially isolated and fragmented by higher-energy collisional dissociation (HCD) at 27% with an injection time of 120 ms, 35000 resolution and target of 2 × 10^5^ counts. An isolation width of 1.8 m/z was applied and underfill ratio was set to 1% and dynamic exclusion to 15 sec.

#### Data analysis

Immunopetidomics data were searched against a custom database for each cell line using PEAKS^®^ X software (Bioinformatics Solutions Inc., Waterloo, ON, Canada) and peptide identities determined applying a false discovery rate cutoff of 1%. The custom database was built by appending RNAseq data from DFT2 and DFT1+IFNγ cells (details of RNAseq provided in supplementary methods) to the reference genome for the Fibroblast cell line, “Salem” (https://www.ensembl.org/Sarcophilus_harrisii/Info/Index). The DFT1 and DFT2 fastq files were aligned with the reference genome using HISAT-2 (hierarchical indexing for spliced alignment of transcripts) (42). Alignments were filtered for a mapping quality greater than 20 and variant call files (VCF) created for a maximum read depth of 50,000 using SAMtools (43). Variants were called using BCFtools (44) and filtered for a minimum read depth of 10 and mapping quality of 20. DFT1 and DFT2 protein databases were then created using R package customProDB as described in (45). The devil reference genome annotation inputs for customProDB were obtained from Ensembl BioMart (46). The non-synonymous SNP and indel variant databases for DFT1 and DFT2 were appended to the devil protein database to create the Sarcophilus_harrisii.DEVIL7.0.custom_db.fasta database that was used for proteomic searches.

### Proteomics analysis

DFT1+IFNγ, DFT2 and Fibroblasts cells (n=3) were lysed in buffer containing 1% sodium deoxycholate (SDC), 7.5 mM TCEP, 50mM TEAB as previously described (47). The lysates were sonicated to sheer the DNA and clarified by centrifugation (16060*xg*, 5 min, RT) before being reduced (60 °C, 15 min in 10 mM TCEP) and alkylated (40 mM chloracetamide, 15 min, RT). The proteins were trypsin digested (ON, 37 °C). SDC was removed by acid precipitation using trifluoroacetic acid followed by centrifugation (16060*xg*, 15 min). The resultant peptides were fractionated using a 10 cm x 4.6 mm Chromolith C18 column running a mobile phase consisting of buffer A (aqueous 0.1% TFA) and buffer B (80% ACN/0.1% TFA). The peptides were eluted using a linear gradient of 5 to 40 % buffer B over 30 min collecting 38 fractions which were concatenated into 7 pools before centrifugal evaporation. Samples were reconstituted using 0.1% formic acid and analysed using mass spectrometry as described above. RefSeq IDs from source proteins were searched on the NCBI RefSeq database to identify the Gene ID for analysis. Proteins which have not been previously annotated in the Tasmanian devil genome (https://www.ensembl.org/Sarcophilus_harrisii/Info/Index) were manually annotated based on protein homology to other species. Due to a lack of annotations for specific protein isoforms in the Tasmanian devil, multiple isoforms were discarded from functional analysis and a single Gene ID was used. The GeneCards database was used to determine which peptide source proteins are overexpressed in the nervous system. Source proteins which are significantly overexpressed in DFT1+IFNγ or DFT2 compared to Fibroblasts were visualised as a Volcano plot in RStudio using the geom_point function of the ggplot package.

### Allotype identification of MHC-I heavy chain fractions

The heavy chains of MHC-I molecules isolated during immunoprecipitation were utilised for allotype identification using mass spectrometry. MHC-I heavy chains from the immunoprecipitation (n=4) were collected and were adjusted to pH 8 followed by reduction with TCEP (12.5 mM, 60 °C, 15 min) and alkylation (25 mM iodoacetamide, 15 min, RT). Proteins were then digested in-solution using trypsin and chymotrypsin (16 h at 37 °C and 25 °C respectively). The resultant peptides were acidified, desalted using Bond-Elut C18 solid phase extractiontips (Agilent) and subjected to mass spectrometry analysis as described above. Data obtained was searched against the Tasmanian devil protein database obtained from NCBI that included the MHC-I sequences using Peaks X (BSI) software, using following settings: fixed modification: carbamidomethyl; variable modifications: phosphorylation (STY), Deamidation (NQ), Oxidation (M); Parent mass error tolerance:10 ppm, fragment mass tolerance 0.02 Da. The peptide allocation to each MHC-I allotype was then manually checked by aligning peptides to the MHC-I alleles defined in (48) to ensure no duplicate assignments were made. Only variants found in at least three out of four replicates were considered for analysis.

### Data analysis and visualisation

Analysis of peptides was performed using a combination of R 3.5.1 & RStudio 1.1.456 and Microsoft Office excel. Some manual annotation of proteins was conducted where necessary. Cellular localization and biological processes of source proteins were analysed using Software Tool for Researching Annotation of Proteins (STRAP, (49)) by inputting Uniprot devil IDs retrieved from Ensemble protein IDs. GO slim name analysis was performed using the Generic GO term Mapper (https://go.princeton.edu/cgi-bin/GOTermMapper), to assign devil gene symbols to human GO slim terms. Ambiguous or unannotated gene symbols were automatically removed by the software. Results from both analyses were compared with six previously published human, mouse and bat MHC peptide datasets: annotations were compiled for three human MHC class I alleles (HLA-A*01:01, HLA-B*27:05, HLA-Cw*04:01 (24, 25, 32)), two murine alleles (H-2D^b^, H-2 K^d^ (26)) and one bat allele (Ptal-N*01:01 (17)), and analyses performed as described above with the exception of murine datasets which were mapped to murine gene symbols.

Consensus-binding motifs were derived from the frequencies calculated for each amino acid at each position along the analysed peptides. Motifs were also analysed using GibbsCluster-2 (https://services.healthtech.dtu.dk/service.php?GibbsCluster-2.0), which clusters peptide sequences into groups, using the algorithm described in (23). Analysis was restricted to “MHC class I ligands of same length” and motif length was adjusted according to analysed sequences (either 8mers or 9mers) and number of clusters changed to 1-6 for all cell lines based on the highest number of MHC-I variants found in the cell lines (Fibroblasts).

The frequency of small and bulky amino acid usage was calculated for each peptide length in the range 7-15mers. The frequencies of leucine at p3 in devil peptides were compared with those of other MHC-I alleles in human and mouse calculated from data available in the online database Immune Epitope Database and Analysis Resource (IEDB). Representative MHC-I alleles were selected from available publications (24–26, 32, 50–53) and only those binding both 8mers and 9mers were considered for comparison.

### Computational analysis of anchor pockets and epitope predictions

Anchor pocket motif prediction for the devil MHC class I (MHC-I) alleles, including those found in the heavy chain fractions from Fibroblasts, DFT1+IFNγ and DFT2 cells, was performed using a custom R-script available at: github.com/sct1g15/MHC_epitope_Prediction_Final. This script aligned 16,748 classical and non-classical alleles across 7 different species to a devil MHC-I query sequence using MAFFT multiple sequence alignment (54). Sequences used for the alignment were taken from the Immuno Polymorphism Database (55). A cut-off point of 0.93*Maximum alignment score was used to determine relevant hits. Anchor pockets/epitopes of devil MHC-I alleles were predicted by identifying sequence homology at the pocket level with experimentally defined peptide epitopes, allowing the inference of similar pocket motifs. This was achieved through implementation of the PMBEC point specific scoring matrix (PSSM), which was derived through competitive assays of combinatorial peptide mixtures to displace MHC-bound and tagged peptide (56). This approach assumes that amino acid modifications to the peptide ligand used to calculate the PSSM would have similar impact on binding affinity when imposed on the receptor sequence. Query sequences were scored through sequence similarity using the PSSM, at the specified pocket positions, which are pre-defined (residues defined below). Each residues contribution to the final alignment score is weighted by its probability to contact the PBR defined through the COACH binding site prediction server following I-TASSER homology modelling of the query sequence (57). This allowed inclusion of adjacent residues enabling appropriate scoring with minor perturbations in pocket sites between MHC-I alleles. Two pre-defined pocket sites were included: B-pocket composed of residues 7, 9, 24, 34, 45, 63, 66, 67, 70, 99 and F-pocket composed of residues 77, 80, 81, 84, 94, 115, 122, 142, 145, 146 as well as residues adjacent to those defined. All positions are set relative to a HLA-A2:01:01:01 reference sequence after multiple sequence alignment. Bat, Chicken and Devil MHC sequences included in the alignment set were not used for epitope prediction.

## Acknowledgements

AB is funded by CRUK Centres Network Accelerator Award A21998 and TE is funded by CRUK Programme Grant A28279. The authors acknowledge the Monash Proteomics & Metabolomics Facility for the provision of mass spectrometry instrumentation and technical support. AWP is supported by NHMRC Principal Research Fellowship 1137739. Computational resources were supported by the R@CMon/Monash Node of the NeCTAR Research Cloud, an initiative of the Australian Government’s Super Science Scheme and the Education Investment Fund.

