## Supplementary Figures for "Passage of transmissible cancers in the Tasmanian devil is due to a dominant, shared peptide motif and a limited repertoire of MHC-I allotypes"

Supplementary figure1

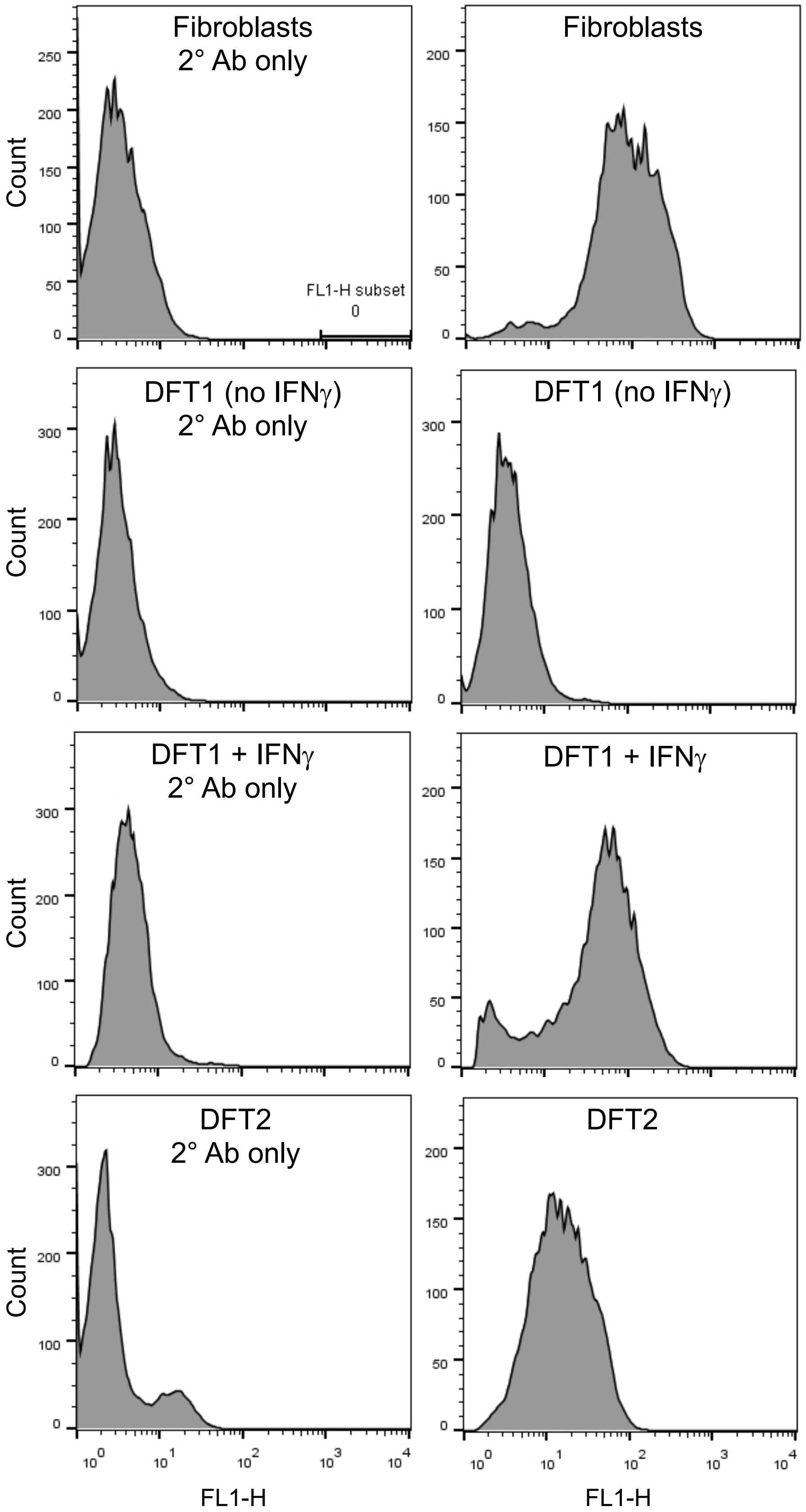


**Supplementary figure 1. Analysis of the expression of MHC class I (MHC-I) molecules in a devil fibroblast cell line and in devil facial tumour disease (DFT1 & DFT2) cell lines.** (a) FACS analysis of surface MHC-I molecules using an antibody against devil B2m in fibroblasts, DFT1 (±IFNy and DFT2 cell lines. 2˚ Ab = negative control using secondary antibody.

Supplementary figure 2

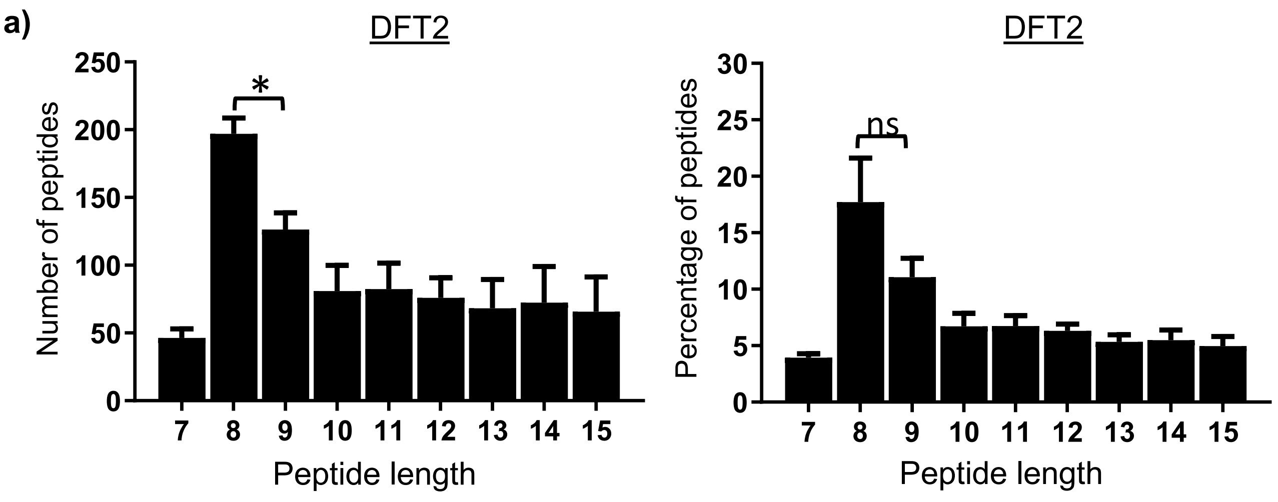


b) **8mers (381)** | **Consensus binding motifs** | **9mers (238)**

| P1 | P2 | P3 | P4 | P5 | P6 | P7 | P8 | | P1 | P2 | P3 | P4 | P5 | P6 | P7 | P8 | P9 |
|---|---|---|---|---|---|---|---|---|---|---|---|---|---|---|---|---|---|
| | | **L** | | | | | **L** | **Dominant** (>30%) | | | **L** | | | | | | |
| **D** | | | | **L** | | | | **Strong** (>20%) | | | | **P** | | | | | **L** |
| E | Q | | I | I | E | E | V | **Moderate** (>10%) | V | R | | L | L | P | V | E | F |
| | R | | R | V | | | | | | Q | | | V | L | | P | |
| | | | L | | | | | | | | | | | | | | |
| | | | P | | | | | | | | | | | | | | |
| | | | K | | | | | | | | | | | | | | |

- Small non-polar (G, A, S, T)
- Hydrophobic (C, V, I, L, P, F, Y, M, W)
- Polar (N, Q, H)
- Negatively charged (D, E)
- Positively charged (K, R)

**Supplementary figure 2. Peptidomics experiments on a smaller number of devil facial tumour 2 (DFT2) cells confirms dominance of 8mer peptide sequences.** (a) Length distribution and (b) motifs of peptides isolated from a smaller number ($1x10^8$) of DFT2 cells (n=3). Both the percentages and total number of peptides for 7 to 15mers are represented as mean±SEM. * = $p<0.05$; ns = not significant. Consensus binding motifs (b) were obtained and represented as described for figure 4; for full dataset see supplementary table 5.

Supplementary figure 3

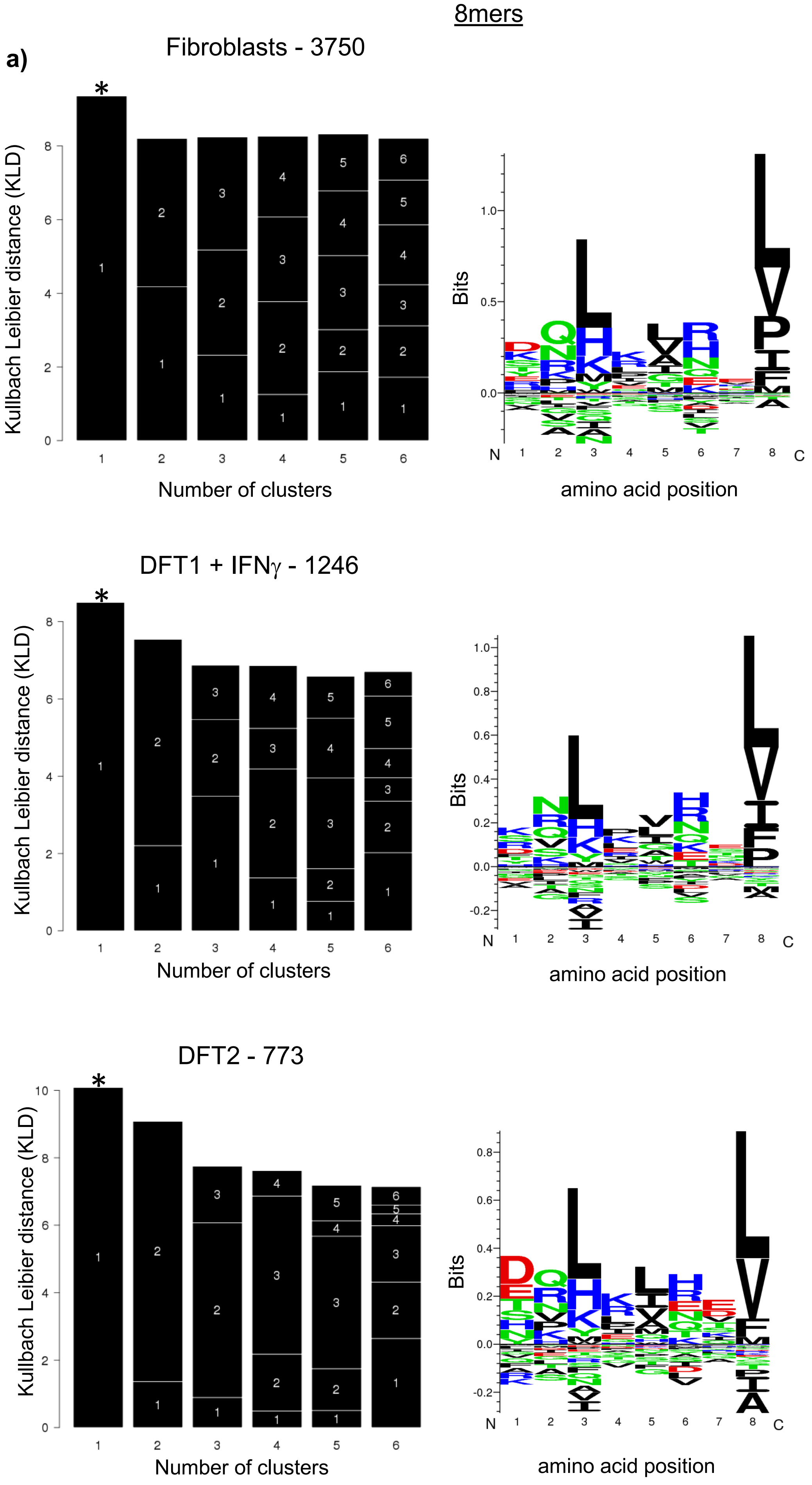


**Supplementary figure 3. Gibbs cluster analysis identifies only one binding motif for 8mers and 9mers from devil fibroblasts and tumour cell lines (DFT1+IFNy & DFT2).** Motifs analysis for 8mers (a) and 9mers (b) performed using GibbsCluster-2, a server for unsupervised analysis and clustering of peptide data (https://services.healthtech.dtu.dk/service.php?GibbsCluster-2.0), which uses as input a list of peptide sequences and attempts to cluster them into meaningful groups. "Number of cluster" on the bar graphs refers to the number of possible motifs, whilst the "Kullbach Leibler Distance (KLD)" gives a measurement of the likelihood that a particular number of motifs (clusters) can be found within that particular peptide pool; the greatest the KLD, the highest the probability of having a certain number of motifs. For each cell line, the most likely number of motifs (greatest KLD, indicated with *) was represented on the left by using logo plots given by GibbsCluster-2; the bigger the size of the letter representing a certain amino acid, the strongest the presence of that amino acid at that position. Numbers after cell line names refer to the number of unique peptide sequences analysed across replicates (n=4).

Supplementary figure 3

9mers

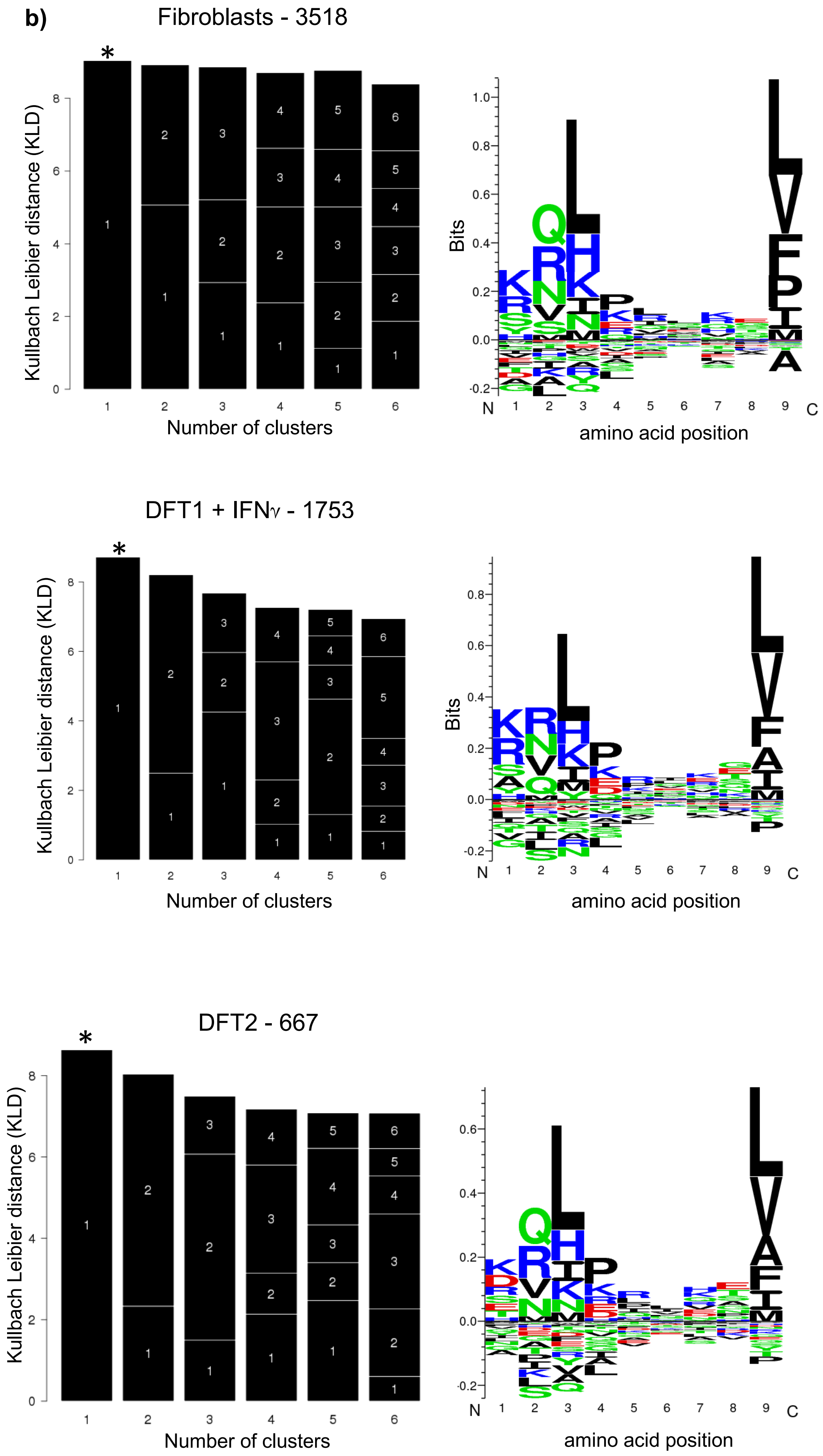

<u>Supplementary figure 4</u>

<u>GO slim name analysis</u>

**a)** Cellular component

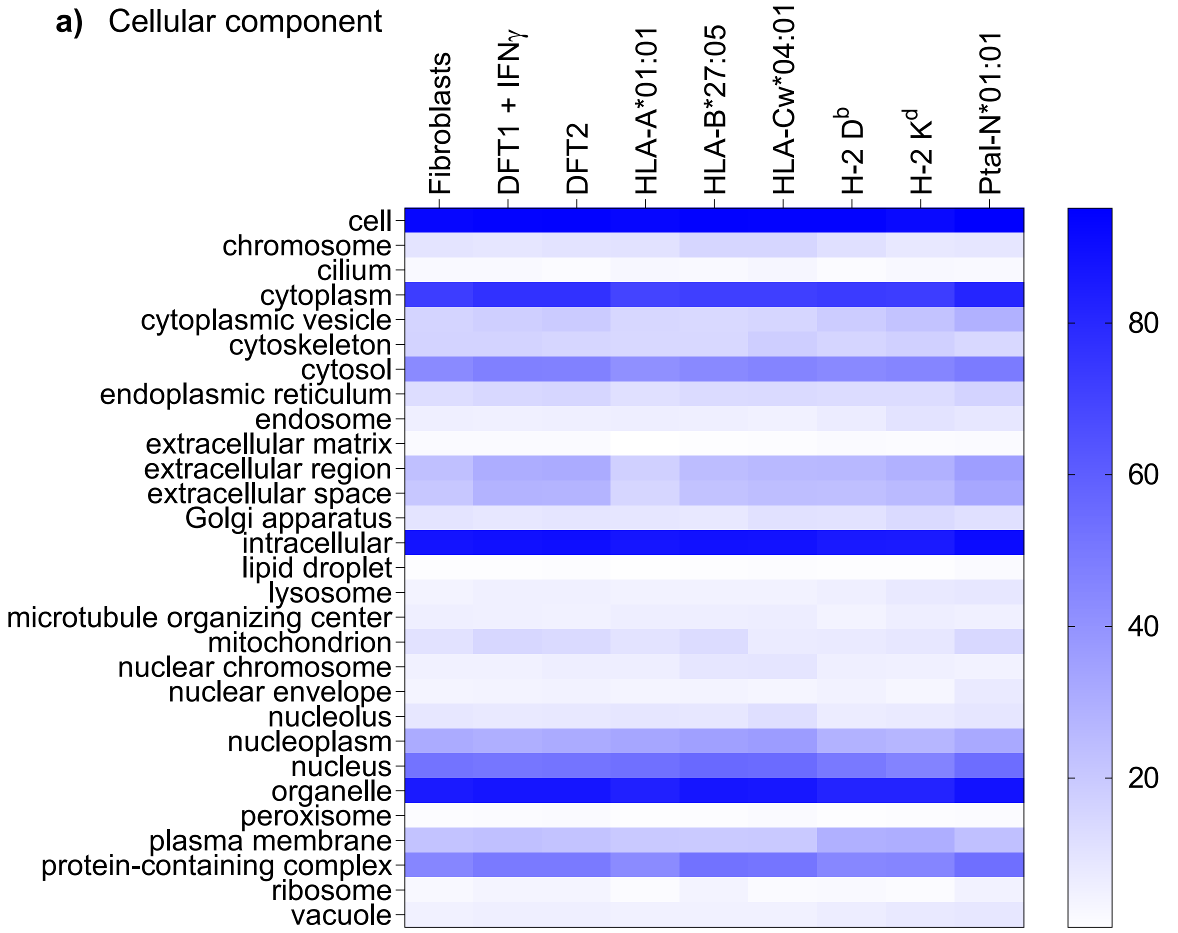


**b)** Biological process

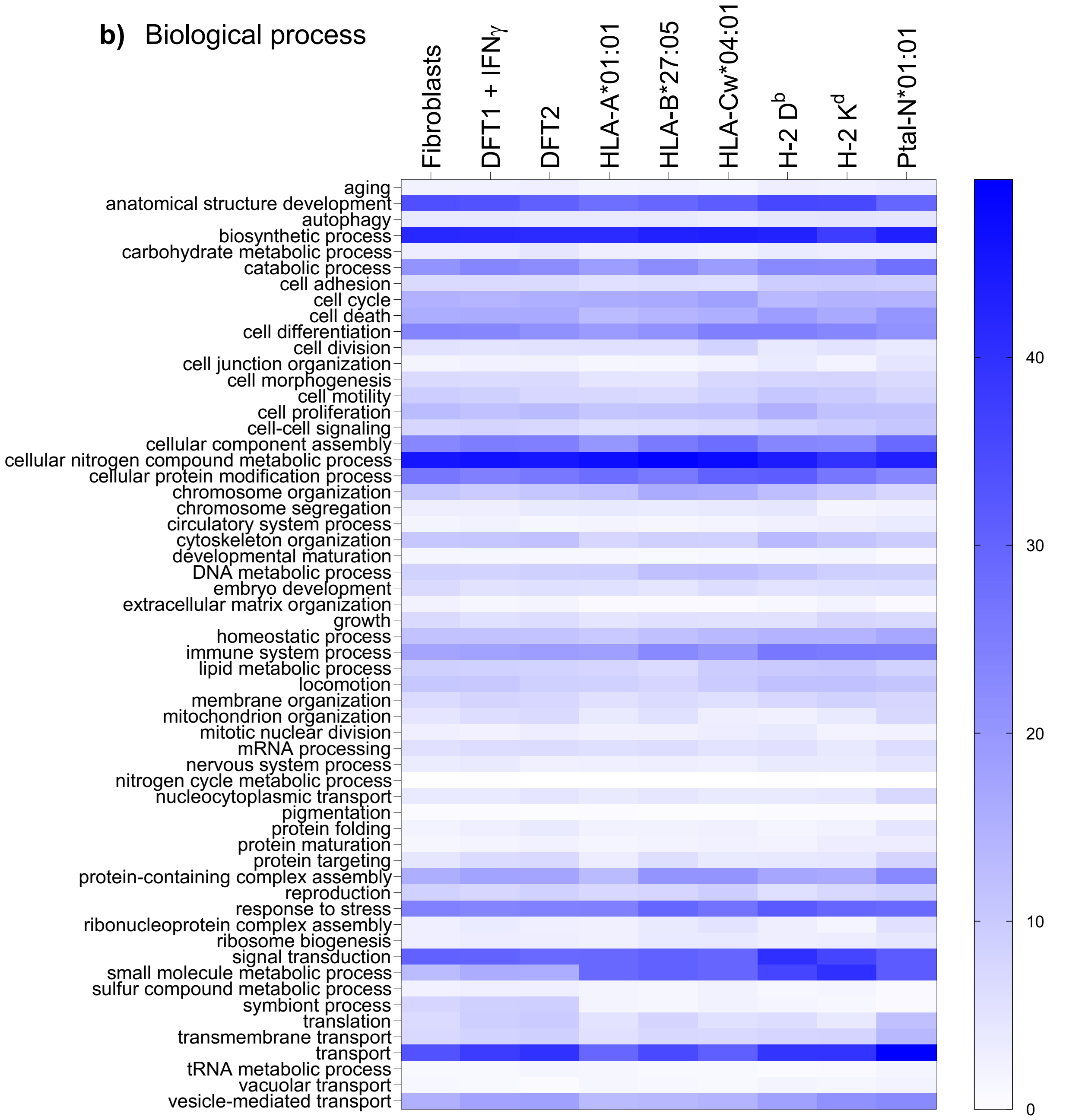

**Supplementary figure 4**. **Source proteins of peptides in host devil and devil facial tumour disease cell lines are similar to those of other species.** Heat maps representing the percentage of source proteins from different cellular components (a) or biological processes (b) for peptides isolated from Fibroblasts, DFT1+IFNy and DFT2 compared with three MHC class I alleles found in human (HLA-A*01:01, HLA-B*27:05, HLA-Cw*04:01), two found in mice (H-2 $D^b$, H-2 $K^d$) and one found in Pteropus alecto (Australian black flying fox, a bat species (Ptal-N*01:01)). Percentages are represented on a sliding scale of the colour blue, were higher percentages corresponds to higher intensity of blue. Gene ontology (GO) slim name analysis using Generic GO term Mapper (https://go.princeton.edu/cgi-bin/GOTermMapper) was performed on unique peptides as described in “methods”. For full dataset see supplementary table 7.
