## Supplementary methods for "Passage of transmissible cancers in the Tasmanian devil is due to a dominant, shared peptide motif and a limited repertoire of MHC-I allotypes"

**Validation of peptide motifs**

Materials and Buffers: Anti-devil β2m antibody was purified by fast protein liquid chromatography (FPLC) from supernatant of hybridoma cells (identifier: 13-34-38). All chemicals were from Sigma-Aldrich unless otherwise stated; all wash buffers and solutions were made using MS grade H_2_O.

Generation of cell lysate: frozen cell pellets (1x10^8^ cells/replicate, n=3) were thawed on ice and resuspended in 5 mL lysis buffer (20 mM TrisCl (pH 8), 150 mM NaCl, 0.5 % (v/v) IGEPAL-630, 0.25 % (w/v) Sodium deoxycholate, 0.2 mM iodoacetamide, 1 mM EDTA, 1x protease inhibitors), disrupted by repeated pipetting and incubated (1 h, 4 °C) with constant rotation. Lysates were centrifuged (2000*xg*, 10 mins, 4 °C) to remove nuclei, and the supernatant was clarified by centrifugation (15000*xg*, 1 h, 4 °C).

Preparation of immunoaffinity columns: 2 mL of protein A sepharose resin (GE healthcare, supplied as a 50 % slurry in 20 % ethanol) was washed (PBS, 10 c.v.) before incubation with anti-devil β2m antibody (2 mg in PBS) under constant rotation (4 °C, 1h). The antibody bound-resin was washed with borate buffer (0.1 M Boric acid, 0.1 M KCl, 4 mM NaOH in, pH 8, 20 c.v.), followed by freshly prepared triethanolamine (0.2 M, pH 8.2; 15 c.v.) to ensure there were no residual amines interfering with the cross-linking reaction. The antibody was cross-linked by incubating (1 h at RT) with DMP (40 mM in 0.2 M triethanolamine, pH 8.2), and the reaction was terminated by adding ice-cold Tris (0.2 M, pH 8, 10 c.v.). Unbound antibody was removed by washing with citrate buffer (0.1 M, pH 3, 10 c.v.) and the column washed with 50 mM Tris (pH 8, 10 c.v.) until pH of flow through was > 7.

Immunoaffinity purification of devil MHC-I molecules: Cross linked antibody-sepharose beads were suspended in 20 mM TrisCl (2mL, pH 8), added to 5 mL of clarified lysate and incubated (16 h, 4 oC) with rotation to capture MHC-I complexes. Lysate-bead slurry was returned to a clean column and the flow through collected and stored at -80 ^o^C for subsequent analysis. The column was then washed at room temperature with the following ice-cold buffers (20 c.v. each): wash buffer 1 (20 mM TrisCl (pH 8), 150 mM NaCl, 0.5 % (v/v) IGEPAL-630, 0.25 % (w/v) Sodium deoxycholate, 0.2 mM iodoacetamide, 1 mM EDTA, 1x protease inhibitors), wash buffer 2 (20 mM Tris (pH 8), 150 mM NaCl), wash buffer 3 (20 mM Tris (pH 8), 400 mM NaCl) and wash buffer 4 (20 mM Tris, pH 8). MHC-I molecules were eluted with 10% acetic acid (5 c.v.). Eluted MHC-I molecules were vacuum dried (RT, overnight) and stored at 4 ^o^C until separation by HPLC.

Separation of MHC-I eluate by RP-HPLC: The dried eluate was re-suspended in 500 $\mu$L of 0.1% Trifluoroacetic acid (TFA)/1% acetonitrile (ACN) was injected using a Thermo Ultimate 3000 HPLC system onto a Chromolith High Resolution RP-18 endcapped 100 mm x 4.6 mm HPLC column (Merck). Peptides were eluted over a linear gradient of 2%-30% buffer B (ACN and 0.1% TFA(v/v)) and fractions collected over 8 min. Fractions were pooled as odd and even fractions, lyophilized and then re-suspended in 20 $\mu$L of water containing 1% Formic acid (v/v) and split into four samples, two odd and two even, for mass spectrometry analysis.

LC-MS/MS analysis of HLA-I peptides: HLA peptides were separated by an Ultimate 3000 RSLC nano system (Thermo Scientific) using a PepMap C18 EASY-Spray LC column, 2 $\mu$m particle size, 75 $\mu$m x 75 cm column (Thermo Scientific) in buffer A (0.1% Formic acid) and coupled on-line to an Orbitrap Fusion Tribrid Mass Spectrometer (Thermo Fisher Scientific, UK) with a nano-electrospray ion source. Peptides were eluted with a linear gradient of 3%-30% buffer B (Acetonitrile and 0.1% Formic acid (v/v)) at a flow rate of 300 nL/min over 110 minutes. Full scans were acquired in the Orbitrap analyser using the Top Speed data dependent mode, preforming a MS scan every 3 second cycle, followed by higher energy collision-induced dissociation (HCD) MS/MS scans. MS spectra were acquired at resolution of 120,000 at 300 m/z, RF lens 60% and an automatic gain control (AGC) ion target value of 4.0e5 for a maximum of 100 ms. MS/MS resolution was 30,000 at 100 m/z. Higher‐energy collisional dissociation (HCD) fragmentation was induced at an energy setting of 28 for peptides with a charge state of 2–4, while singly charged peptides were fragmented at an energy setting of 32 at lower priority. Fragments were analysed in the Orbitrap at 30,000 resolution. Fragmented m/z values were dynamically excluded for 30 seconds.

Database searching: Analysis of raw data was performed using Peaks Ver 8.5 software (Bioinformatics Solutions, Inc, Canada) and the data processed to generate reduced charge state and deisotoped precursor and associated product ion peak lists. These peak lists were searched against a custom Sarcophilus_harrisii database: Sarcophilus_harrisii.DEVIL7.0.custom_db.fasta (30/11/2017, 24,670 entries) using a non-specific search. Peaks PTM searches were performed using carboxyamidomethylation of cysteine as a fixed modification and variable modifications were set to contain oxidation of methionine. The false discovery rate (FDR) was estimated with randomized decoy database searches and were filtered to 1% FDR.

**DFT1 and DFT2 transcriptomes**

DFT2 cells (red velvet/DFT2_202) and DFT1 cells (4906) were cultured and DFT1 cells were treated with IFNy as described in the main methods. Cells were harvested and mRNA was extracted using the Nucleospin RNA mini kit (Machenery and Nagel) and following the manufacturers instructions. The quality of RNA was determined using the Agilent 2100 Bioanalyzer. 1000ng of RNA was used to generate strand specific RNA libraries. Libraries were sequenced on an Illumina Hiseq by Eurofins Genomics, generating over 64 million 150bp paired end reads for each cell line. Data analysis is described in the main methods.
